# Callosal anisotropy predicts attentional network changes after parietal inhibitory stimulation

**DOI:** 10.1101/2020.07.13.196824

**Authors:** Selene Schintu, Catherine A. Cunningham, Michael Freedberg, Paul Taylor, Stephen J. Gotts, Sarah Shomstein, Eric M. Wassermann

**Author notes:** **Corresponding author**: Selene Schintu, Ph.D., Behavioral Neurology Unit, National Institute of Neurological Disorders and Stroke, Building 10, Room 7D48, 10 Center Drive, MSC 1440, Bethesda, MD 20892-1430.

## Abstract

Hemispatial neglect is thought to result from disruption of interhemispheric equilibrium. Right hemisphere lesions deactivate the right frontoparietal network and hyperactivate the left via release from interhemispheric inhibition. Support for this theory comes from neuropsychological evidence as well as transcranial magnetic stimulation (TMS) studies in healthy subjects, in whom right posterior parietal cortex (PPC) inhibition causes neglect-like, rightward, visuospatial bias. Concurrent TMS and fMRI after right PPC TMS show task-dependent changes but may fail to identify effects of stimulation in areas not directly activated by the specific task, complicating interpretations. We used resting-state functional connectivity (RSFC) after inhibitory TMS over the right PPC to examine changes in the networks underlying visuospatial attention.

In a crossover experiment in healthy individuals, we delivered continuous theta burst TMS to the right PPC and vertex as control condition. We hypothesized that PPC inhibitory stimulation would cause a rightward visuospatial bias, decrease PPC connectivity with frontal areas, and increase PPC connectivity with the attentional network in the left hemisphere. We also expected that individual differences in fractional anisotropy (FA) in white matter connections between the PPCs would account for variability in TMS-induced RSFC changes.

As expected, TMS over the right PPC caused a rightward shift in line bisection judgment and increased RSFC between the right PPC and the left superior temporal gyrus. This effect was inversely related to FA in the posterior corpus callosum. Local inhibition of the right PPC reshapes connectivity in the attentional network and depends on interhemispheric connections.

## 1. INTRODUCTION

The parietal cortex is the hub of the attentional network (Behrmann et al., 2004). Its central role is supported by the literature on neglect patients, in whom a lesion to the right parietal cortex produces loss of awareness of the contralesional side of space (Critchley, 1953; Vallar, 1998; Mort et al., 2003). According to the interhemispheric rivalry theory (Kinsbourne, 1977), hemispatial neglect results from breakdown of the interhemispheric balance normally maintained by reciprocal interhemispheric inhibition. Right parietal lesions not only decrease local activation but increase activity in the contralateral hemisphere (Corbetta et al., 2005) by releasing it from inhibition. During spontaneous recovery from neglect, a “push-pull” pattern of right-side re-activation and left-side de-activation has been observed in the posterior parietal cortex (PPC; Corbetta et al., 2005).

Additional support for the interhemispheric rivalry theory of neglect comes from transcranial magnetic stimulation (TMS) studies. Nominally inhibitory TMS applied over the left PPC reduces neglect (for review see Cazzoli et al., 2010). Conversely, when inhibitory TMS is applied to the right PPC in healthy individuals, it causes a temporary, neglect-like, rightward visuospatial bias (Fierro et al., 2000; Hilgetag et al., 2001; Bjoertomt et al., 2002; Hung et al., 2005; Dambeck et al., 2006; Sack et al., 2007; Nyffeler et al., 2008; Cazzoli et al., 2009). Koch et al. (2008) found evidence that the connection from the left PPC to the ipsilateral primary motor area (M1) is hyperexcitable in neglect patients, by delivering conditioning stimulation to the PPC and testing the motor evoked potential from M1 stimulation a few ms later. The PPC-M1 connection is part of the reciprocal frontoparietal circuit critical for visuospatial attention (Doricchi and Tomaiuolo, 2003; Bartolomeo et al., 2007; He et al., 2007; Thiebaut de Schotten et al., 2011) and excitability in the PPC-M1 pathway is considered a proxy for the state of the greater frontoparietal attentional network in that hemisphere. However, direct evidence for the ability of left PPC TMS to modulate the functional connectivity of the frontoparietal network consists of a single case study (Bonnì et al., 2013). TMS studies with only behavioral outcomes can support mechanistic theories but do not provide direct evidence that the targeted network has been actually engaged. The motor evoked potential is well understood and easily quantified, but it is generated by the corticospinal motor output system whose relationship with the frontoparietal network controlling attention is not understood.

Functional magnetic resonance imaging (fMRI) coupled with TMS can partially overcome this limitation. Concurrent TMS/fMRI paradigms measuring task-dependent activation after right PPC inhibitory TMS have found decreased activity in ipsilateral nodes of the frontoparietal network (Sack et al., 2007; Ricci et al., 2012) and increased activation in contralateral regions, such as the primary visual area (Heinen et al., 2011). Using fMRI, Sack et al. (2017) found that the stimulated area was functionally connected to the frontoparietal network bilaterally, but only during task execution. However, task-related activation may fail to identify areas whose connectivity is affected by the stimulation but are not activated by the specific task. Studies of resting-state functional connectivity (RSFC) in the attentional network after PPC TMS can provide information on the network independent of activation and measure the baseline state of the network affected in neglect.

We used RSFC to study the network effects of nominally inhibitory right PPC TMS in healthy participants. Based on the evidence above, we hypothesized that inhibition of the right PPC would result in a rightward visuospatial bias with a corresponding decrease of PPC connectivity with frontal areas and/or increase in connectivity with contralateral areas of the attentional network.

We expected that any contralateral changes would depend on anatomical connectivity between two PPCs. The PPCs are connected by fibers in the posterior part of corpus callosum (Koch et al., 2011), and micro- and macrostructural differences in this pathway account for significant interindividual variability in the attentional shift produced by PPC inhibitory TMS (Chechlacz et al., 2015). We therefore hypothesized that interindividual differences in fractional anisotropy (FA) in this pathway would partially predict individual RSFC changes induced by inhibitory TMS over the right PPC.

## 2. MATERIALS AND METHODS

The study was pre-registered at https://aspredicted.org/blind.php?x=2gx7bk

### 2.1 Participants

Seventeen adults (11 female; age = 25.94 ± 1.01SEM), free of neurological disorders or medications affecting brain function, participated in the study. All had normal or corrected-to-normal vision, were right-handed (Edinburgh Inventory; Oldfield, 1971), and were right-eye dominant (hole-in-card test; Miles, 1930). Participants were compensated and gave written informed consent. The study was approved by the National Institutes of Health, Central Nervous System Institutional Review Board, and conducted in accordance with the ethical standards of the 1964 Declaration of Helsinki (World Medical Association, 2013).

### 2.2 Procedure

The crossover-design experiment consisted of two sessions of behavioral testing and fMRI, one before and one after TMS (Figure 1). In each session participants underwent resting state and anatomical scans. Following the first resting state scan, participants carried out the perceptual and manual line bisection tasks along with two tasks assessing spatial bias in proprioceptive (straight-ahead pointing) and sensorimotor (open-loop pointing) performance (Figure1). Participants then received TMS over either the right PPC or the vertex in counterbalanced order. Immediately after TMS, participants performed the straight-ahead and open-loop pointing tasks (early post-adaptation), underwent the second resting state scan, and performed the perceptual and manual line bisection, straight-ahead and open-loop pointing tasks again (late post-adaptation). Diffusion weighted imaging (DWI) data were acquired in a separate session (Figure 1).

**Figure 1.**
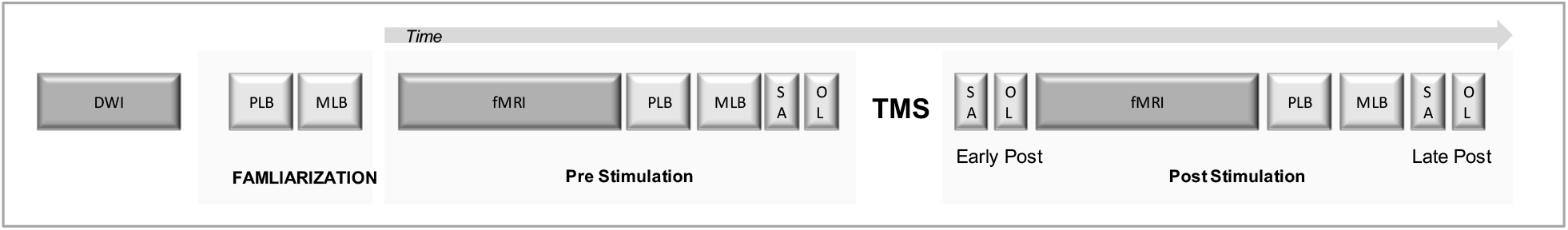
Experimental timeline. DWI = Diffusion Weighted Imaging; PLB = perceptual line bisection; MLB = manual line bisection; SA = straight-ahead pointing; OL = open-loop pointing; fMRI = functional magnetic resonance imaging; TMS = transcranial magnetic stimulation.

During the tasks, participants were seated in front of a horizontal board with their heads supported by a chinrest. On the board, a target (8 mm in diameter) was positioned at 0 degrees from the body midline, at approximately 57cm distance from participant’s nasion and was used for open-loop and straight-ahead pointing tasks.

### 2.3. Behavioral assessment

We used the four behavioral tasks described previously in our RSFC study of prism adaptation (Schintu et al., 2020b).

#### Perceptual line bisection

prioritizes the perceptual, and minimizes the motor, component of the visuospatial bias by asking participants to judge a series of pre-bisected lines instead of actively bisecting them. We used a modified version of the Landmark task (Milner et al., 1992). The task consisted of 66 white, pre-bisected, lines (350 mm × ~2 mm) displayed on a black screen positioned 35 cm from the eyes. Lines were transected at the true center and at 2, 4, 6, 8, and 10 mm to the left and right of the true center. Each of the 11 different pre-bisected lines was presented six times in a pseudorandom order, yielding a total of 66 trials, which took approximately three minutes to complete. Each line was displayed for a maximum of five seconds, or until a response was made, and was then replaced by a black-and-white, patterned mask, which stayed on the screen for one second before the next line was displayed. We used Presentation software (Neurobehavioral Systems, Inc., Berkeley, USA) to generate the stimuli, record responses, and control the task. Participants were instructed to inspect each line and judge whether the transecting mark was closer to the left or right end and to respond within 5 seconds by pressing pedals positioned under the left and right feet. We chose a pedal response to limit the use of the right hand, which was used for PA, since post-adaptation feedback from that hand could contribute to de-adaptation. Participants performed at least ten practice trials before the baseline measurement. For each participant, we plotted the percentage of right-side responses as a function of the position of the transector (true center and 2,4,6,8,10 mm to left and to the right of the true center). We then fit a sigmoid function to the data. The value on the x-axis corresponding to the point at which the participant responded with the right pedal 50% of the time was taken as the point of subjective equality (PSE).

#### Manual line bisection

emphasizes the motor over the perceptual component of the visuospatial bias (Milner et al., 1992). We used this task (Schenkenberg et al., 1980) to measure the visuospatial shift induced by TMS. It consisted of a series of 10 black lines (identical in size to those used for the perceptual line bisection task) drawn on 297mm × 420mm sheets of paper, which were positioned over the computer screen used for the perceptual line bisection task. Participants were instructed to inspect each line and, with a pen held in their right hand, draw a vertical mark at the perceived center of each line. No time limit was imposed, and participants took on average 1 second to place the mark on one line. We measured the distance between the mark placed by the participant and the true center of the line and took the average as the PSE. Marks to the right of center were coded as positive.

#### Straight-ahead pointing

was used to measure the proprioceptive shift induced by TMS. Participants performed six pointing movements to the midline with the right index finger at a comfortable and uniform speed, while resting their left hands on their laps. Before each movement, participants were told to close their eyes, imagine a line splitting their body in half, and project it onto the board in front of them. We then asked them to point to the projected midline with their eyes closed and then return to the starting position when told by the experimenter. To ensure that participants had no visual feedback, the arm and hand were occluded by a cardboard baffle before movement onset. The proprioceptive shift was measured as the average distance between the landing position and the true midline with precision of +/− 0.5 cm.

#### Open-loop pointing

was used to measure the sensorimotor shift induced by TMS. Participants performed six pointing movements with the right index finger to the central (0°) target at a comfortable and uniform speed, while resting their left hands on their laps. Before each movement, we instructed participants to look at the central target, close their eyes, point to the target while keeping their eyes closed, and then return the hand to the starting position when prompted by the experimenter. As in the straight-ahead task, vision of the arm and hand was occluded. The sensorimotor shift was measured as the average distance between the landing position and the central target with a precision of +/− 0.5 cm.

### 2.4 MRI

#### MRI acquisition

Functional and structural MRI data were acquired with a 32-channel head coil in 3-Tesla GE medical systems discovery MR750 scanner. Head movement was minimized with padding. A whole-brain T1-weighted anatomical image (MPRAGE) was obtained for each participant (172 slices, voxel size 1.0 × 1.0 × 1.0 mm^3^, repetition time (TR) = 7.66 ms, echo time (TE) = 3.476 ms, inversion time (TI) = 1100 ms, field of view (FOV) =256 × 156 × 176 mm^3^, flip angle = 7°). T2* blood oxygen level-dependent (BOLD) resting state scans were acquired for all participants (45 slices aligned to the AC-PC axis, voxel size 3.0 × 3.0 × 3.0 mm^3^, TR = 2500 ms, TE = 30.0 ms, FOV = 216 × 135 × 216 mm^3^, flip angle 70°, 72 × 45 × 72 acquisition matrix). During resting state scans, lighting was dimmed, and participants were instructed to lie still, look at a white central cross appearing on a black screen and to think about nothing.

DWI scans were performed using a 32-channel head coil in a 3 Tesla Siemens Magnetom Prisma scanner. Head movement was minimized with padding. Data were acquired using an echo-planer imaging (EPI) sequence with a voxel size of 2 × 2 × 2 mm^3^ and the following parameters: TR = 9200 ms, TE = 79 ms, and matrix size = 110 × 110 × 80. Diffusion imaging was performed in multiple shells, with 70 total diffusion-weighted volumes (10 × b=300 s/mm^2^, 60 × b=1100 s/mm^2^) and 10 non-diffusion-weighted volumes (b=0 s/mm^2^). Two sets of images, with opposite phase-encoded directions (anterior-posterior), were acquired to correct for susceptibility-induced distortion. As anatomical references, high resolution T2-weighted fat-saturated images were acquired with the following parameters: voxel size = 1.1 × 1.1 × 1.7 mm^3^, matrix size = 192 × 192 × 94.

#### MRI preprocessing

We preprocessed the functional and structural MRI data with the AFNI software package (v. 18.2.15; Cox, 1996) and followed the general approach of Schintu et al. (2020b). The anatomical scans were segmented into tissue compartments using the FreeSurfer image analysis suite (http://surfer.nmr.mgh.harvard.edu/). We removed the two initial volumes from each resting state scan to allow the magnetic field to stabilize. Volumes were then despiked to minimize outlying time points in each voxel and slice-time corrected to the first slice. Subsequent to this, co-registration of individual EPI volumes to each other and to the anatomical scan and transformation to standard space (Talairach & Tournoux, 1988) at a 2-mm isotropic resolution was accomplished in a single step so that the data were re-gridded only once. Data were then smoothed with an isometric 4-mm full-width half-maximum Gaussian kernel, and then scaled each voxel time series to a mean of 100, with a range of 0-200. AFNI’s 3dTproject was then used to simultaneously censor TRs with head movements > 0.3 mm, bandpass filter the times series between 0.01 and 0.1 Hz and regress the 6 head motion parameters and their first derivatives (Lindquist et al., 2019). Measures of mean framewise displacement, using the AFNI function @1dDiffMag) and average voxelwise signal amplitude (standard deviation), were also calculated for use as nuisance covariates in group-level analyses to control for residual global artifacts in the resting-state scans (Gotts et al., 2020).

DWI processing was performed using a combination of AFNI’s FATCAT programs (Taylor and Saad, 2003) and the TORTOISE software package (v. 3.1.1). Using FATCAT, the T2-weighted anatomical volumes were axialized (rigidly aligned with reference volumes) to provide standard viewing planes (fat_proc_axialize_anat). Additionally, each set of raw DWIs was visually inspected and any volumes with noticeable dropout or motion artifacts was removed from further processing (@djunct_dwi_selector.tcsh and fat_proc_select_vols). TORTOISE’s DIFFPREP was used on each set of DWIs to reduce effects of subject motion and eddy current distortion (default settings and options used), and then DR_BUDDI was run to correct for B0 inhomogeneities, resulting in a single set of DWIs aligned to the T2-weighted reference for each subject (default options, with distortion level=medium and final resampling to 1.5 mm isotropic voxels). Nonlinear tensor fitting was performed in AFNI (fat_proc_dwi_to_dt), as well as standard RGB-colorized DEC maps of the tensors (fat_proc_decmap), and FreeSurfer results were transformed using linear affine alignment with AFNI to map the parcellations into the final diffusion space (fat_proc_map_to_dti). At each processing step, results were checked visually using quality control (QC) images that are automatically generated for each of the above AFNI commands, as well as for the TORTOISE commands if AFNI is installed.

#### Analysis

##### RSFC

To initiate the analysis, we created a seed region for each participant composed of a 3mm radius sphere around the coordinates of the stimulation target in the PPC (Figure 2). The seed voxel time series were first averaged and then correlated (Pearson) with the time series in each voxel throughout the brain, with the correlations transformed by Fisher’s z to improve normality. After fMRI preprocessing, we created a mask for each participant, including voxels with functional data. For the group analyses, we created a group-level brain mask, using voxels where at least 90% of participants had data.

**Figure 2.**
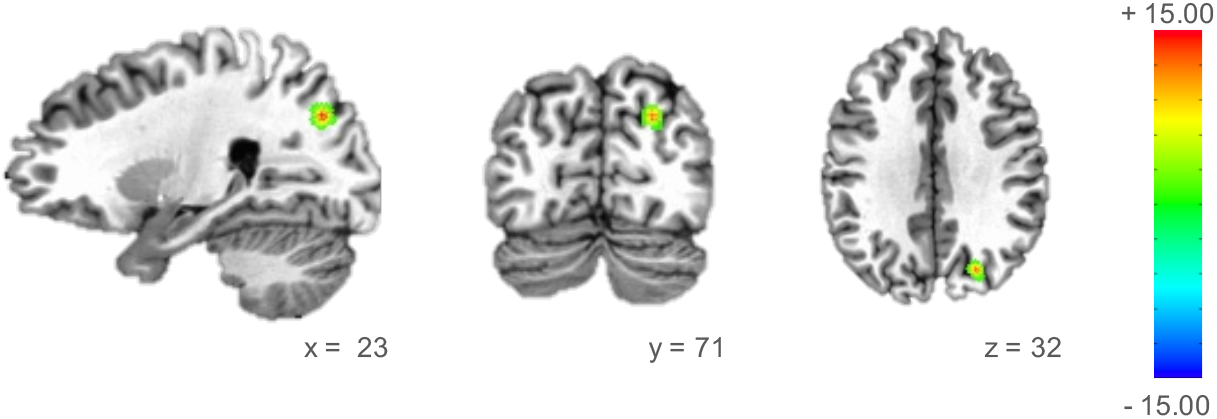
Overlap of the 17 participants’ PPC TMS targets. Color scale indicates number of participants.

##### Diffusion tensor imaging (DTI)

To identify region of interests (ROIs) for the right and left PPC and frontal eye field (FEF), which is the principal frontal node of the dorsal attentional network (Corbetta and Shulman, 2011), we used a probabilistic atlas of the visual areas (Wang et al., 2015). We located ROIs by transforming the individual combined intraparietal sulcus areas 1 and 2 (IPS1/2) and FEF maps into TT_N27 template space, keeping voxels that had ≥ 30% probability of being in IPS1/2 and FEF (Figure 3a). Minimal inflation was applied to all ROIs, informed by the white matter skeleton, to enable tracking. Transcallosal tracts between the right and left IPS1/2, and FEF were reconstructed using full probabilistic tractography in 3dTrackID (Taylor and Saad, 2013; Figure 3b).

**Figure 3.**
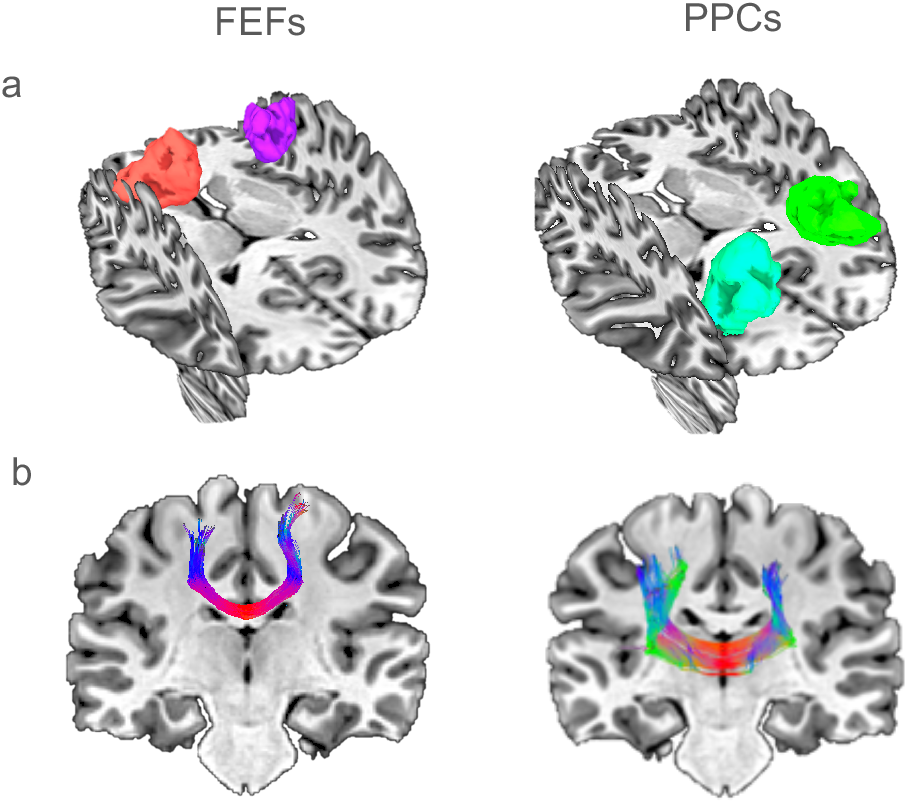
**a.** ROIs used to reconstruct transcallosal tracts between left and right IPS1/2 and FEFs. **b.** Reconstructed tracts between the right and left IPS1/2 and FEF, using full probabilistic tractography in 3dTrackID.

### 2.5 TMS

TMS was delivered with a Magstim Rapid stimulator connected to a Figure-of-eight coil with a diameter of 70 mm. Continuous theta-burst stimulation (3-pulse bursts at 50 Hz delivered every 200 ms; Huang et al., 2005) was delivered at 80% of the active motor threshold (AMT; see below). Stimulation lasted 40 sec with a total of 600 pulses. The electromyogram (EMG) was recorded from the left first dorsal interosseous (FDI) muscle. After finding the scalp location where stimulation evoked the largest motor evoked potential (MEP) from the left FDI, we determined the AMT, defined as the minimum stimulus intensity to produce a MEP > 0.25 mv on 10 consecutive trials, during contraction of the left FDI at approximately 10% of maximum voluntary contraction. Effort was kept constant across participants and over time by monitoring the rectified EMG online.

We used the MRI-guided, Brainsight frameless stereotaxic system to record the optimal scalp location for activating the FDI and locate the TMS target. To identify the PPC target we used a probabilistic atlas of the visual areas (Wang et al., 2015) to find the right intraparietal sulcus area 1 (IPS1). We created a visual target for aiming the coil by transforming the IPS1 map from the probabilistic atlas into TT_N27 template space, using voxels with ≥ 30% probability of being in IPS1. We projected a mask of this region into the subject space and targeted the cortical part of the mask. We held the coil tangential to the scalp and rotated the junction of the windings perpendicular to the closest gyrus (Figure 3). For the vertex condition, identical TMS was applied over the vertex with intersection of the windings in the sagittal plane.

### 2.7 Statistical analysis

The required sample size was estimated using SAS (version 9.4), based on the mean and standard deviation of change in frontoparietal RSFC from a previous study investigating changes in RSFC following motor learning (Albert et al., 2009). Seventeen participants were required to achieve 80% power to reject the null hypothesis at a significance level of 0.05, two-tailed.

Statistical analyses were performed with SPSS (Version 24.0), Matlab (version R2016a) and AFNI (3dLME command; Cox, 1996) with family-wise alpha set at .05. All data are presented as mean and standard error of the mean (SEM). Effect sizes were computed as Cohen’s d. When sphericity was violated, Greenhouse-Geissser corrected values are reported. We used two-tailed paired t-tests for post-hoc comparisons.

To assess the changes in RSFC, data were submitted to a linear mixed effects regression model (LMER using AFNI 3dLME function) with seed-based functional connectivity (correlation maps) as the dependent variable, Time (pre, post), and Stimulation Location (PPC, vertex) as fixed effects, Subject as a random effect, and motion (@1dDiffMag) and average voxel-wise standard deviation as nuisance covariates, cluster size correction p < 0.001 k= 53 (acf-version 3dClustSim; Cox et al., 2017).

## 3. RESULTS

### 3.1 Behavior

#### Perceptual line bisection

A repeated measures ANOVA with Time (pre, post) and Stimulation Location (PPC, vertex) as within-participants variables revealed a significant Time by Stimulation Location interaction [F(1, 14) = 6.084, p = 0.027, *η^2^_p_*=0.303; Figure 4a]. Post-hoc comparison showed that when TMS was delivered over the PPC the PSE shifted significantly rightward from pre (−0.73 mm) to post [−0.07 mm; t(14) = −2.171, p = 0.048, Cohen’s *d* = 0.560] but not when TMS was delivered over the vertex [t(14) = 0.739, p = 0.472, Cohen’s *d* = 0.190]. All other effects were non-significant (all F ≤ 0.299, p ≥ .593).

**Figure 4.**
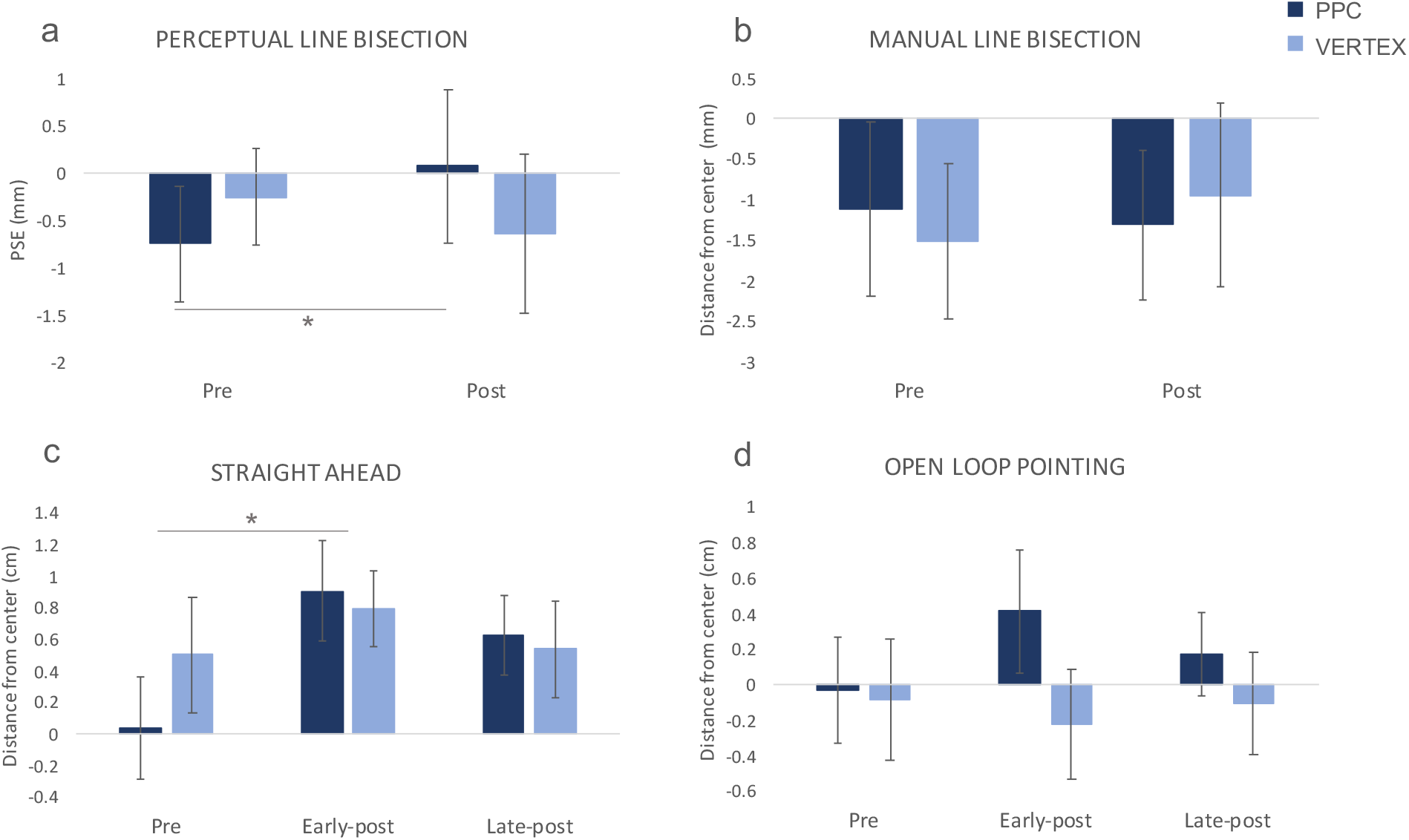
Behavioral effects of TMS. Negative and positive values represent left and right of center, respectively. Error bars represent 1 SEM. * *p < 0.05.*

#### Manual line bisection

Repeated measures ANOVA with Time (pre, post) and Stimulation Location (PPC, vertex) as within-participants variables showed no significant main effect or interaction (all F ≤ .507, p ≥ .487; Figure 4b).

#### Straight-ahead pointing

We measured proprioceptive performance by quantifying the deviation in the pointing between the perceived and true participants’ midline. Repeated measures ANOVA with Time (pre, early-post, late-post) and Stimulation Location (PPC, vertex) as within-participant variables revealed a significant main effect of Time [F(2, 28) = 4.687, p = 0.018, *η^2^_p_*= 0.251; Figure 4c]. Post-hoc comparison revealed that, independently of the Stimulation Location, the pointing error shifted significantly rightward from pre (0.27 cm) to early-post [0.85 cm; t (14) = −2.993, p = 0.020 Bonferroni corrected, Cohen’s *d* = 0.772] but not to late-post [0.58 cm; t (14) = −1.458, p = .334 Bonferroni corrected, Cohen’s *d* = 0.376]. All other effects where non-significant (all F ≤ 1.090, p ≥ .350).

#### Open-loop pointing

We measured sensorimotor performance by quantifying the deviation between the landing position and the true center. Repeated measures ANOVA with Time (pre, early-post, late-post) and Stimulation Location (PPC, vertex) as within-participant variables revealed no significant main effect or interaction (all F ≤ 3.717, p ≥ .074; Figure 4d).

### 3.2 RSFC

#### Seed-based analyses

Before covarying head motion and the average standard deviation nuisance measures in the LMER, we checked for significant baseline differences in the nuisance variables between the PPC and vertex stimulation sessions with a series of two-tailed paired t-tests. We found no differences at baseline [motion t(16) = 1.301, p = .212; standard deviation t(16) = .211, p = .835; PPC: motion mean = .064 (SEM = .006), standard deviation = 1.149 (0.048); vertex motion = .056 (.006), standard deviation = 1.137 (0.054))], or at post [(motion t(16) = −.219, p = .829; standard deviation t(16) = −.949, p = .357; PPC: motion = .071 (.010), standard deviation = 1.162 (.068); vertex: motion .073 (.008), standard deviation = 1.229 (.058)]. Similarly, there was no significant difference between pre- and post-phases for both the PPC [motion t(16) = .953, p = .355 deviation t(16) = .297, p = .770] and vertex [motion t(16) = 2.040, p = .058; standard deviation t(19) = 1.483, p = .172].

The LMER analysis revealed a significant Time × Stimulation Location interaction (p = 0.001, corrected), such that RSFC was differentially affected according to the stimulation site. The analysis detected the left superior temporal gyrus (STG; BA41; 59 voxels/ 177 mm^3^; x = −37 y = −31 z = 16; Figure 4a) as significant cluster relative to the right IPS1 seed.

To follow up the Time x Stimulation Location interaction for each stimulation location, we extracted the time series of the cluster which survived the LMER and computed the correlation between the cluster and IPS1 timeseries, averaged the time series of all voxels in this cluster, and then compared the Fisher z-transformed correlation coefficient before and after TMS. Post-hoc testing of the correlation coefficients revealed that PPC TMS increased RSFC between the right IPS1 and left STG seeds [t(16) = 2.989, p = 0.009 Cohen’s d = 0.72; pre = 0.15, post = 0.26], whereas vertex TMS decreased RSFC between the right IPS1 and left STG [t(16) = −3.331, p = 0.004 Cohen’s d = 0.80; pre = 0.28, post = 0.14] (Figure 5).

**Figure 5.**
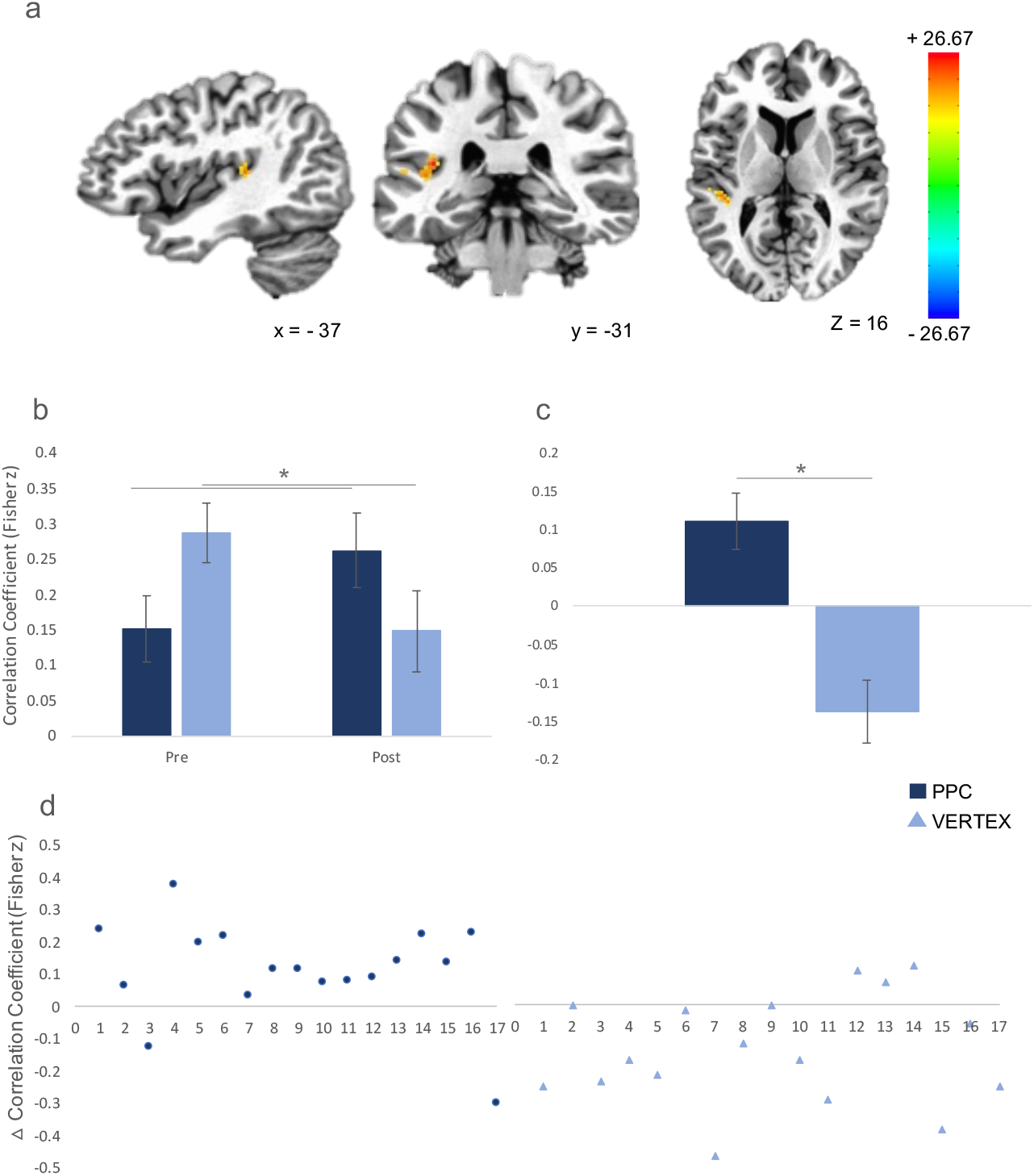
RSFC seed based analysis. **a.** Significant clusters of RSFC change from the Time x Stimulation Location interaction. Color scale indicates F-value. **b.** RSFC between left STG and the right IPS1/2 seed. **c.** Amount of change (post – pre) in RSFC between the left STG and the IPS1/2 seed Error bars = 1 SEM. *p < .05. **d.** Individual data from **c**.

To rule out the possibility that our results were driven by baseline differences between the two stimulation conditions, we performed a repeated measures ANOVA with the change in RSFC (post – pre) as the independent variable, Stimulation Location as the within participants variable and baseline RSFC as a covariate. The results still showed a significant main effect of Stimulation Location [F(1, 14) = 9.817 p = 0.007, *η^2^_p_* = 0.412].

Interindividual variance in the change in RSFC between the right IPS1 and left STG was significantly influenced by FA between the two PPCs when TMS was delivered over the right IPS1 (r = – 0.539, p = 0.026; Figure 6), but not by FA between the FEFs (r = −.279, p = .278). Callosal FA between PPCs and FEFs were strongly correlated within individuals (r = .631, p = .007).

**Figure 6.**
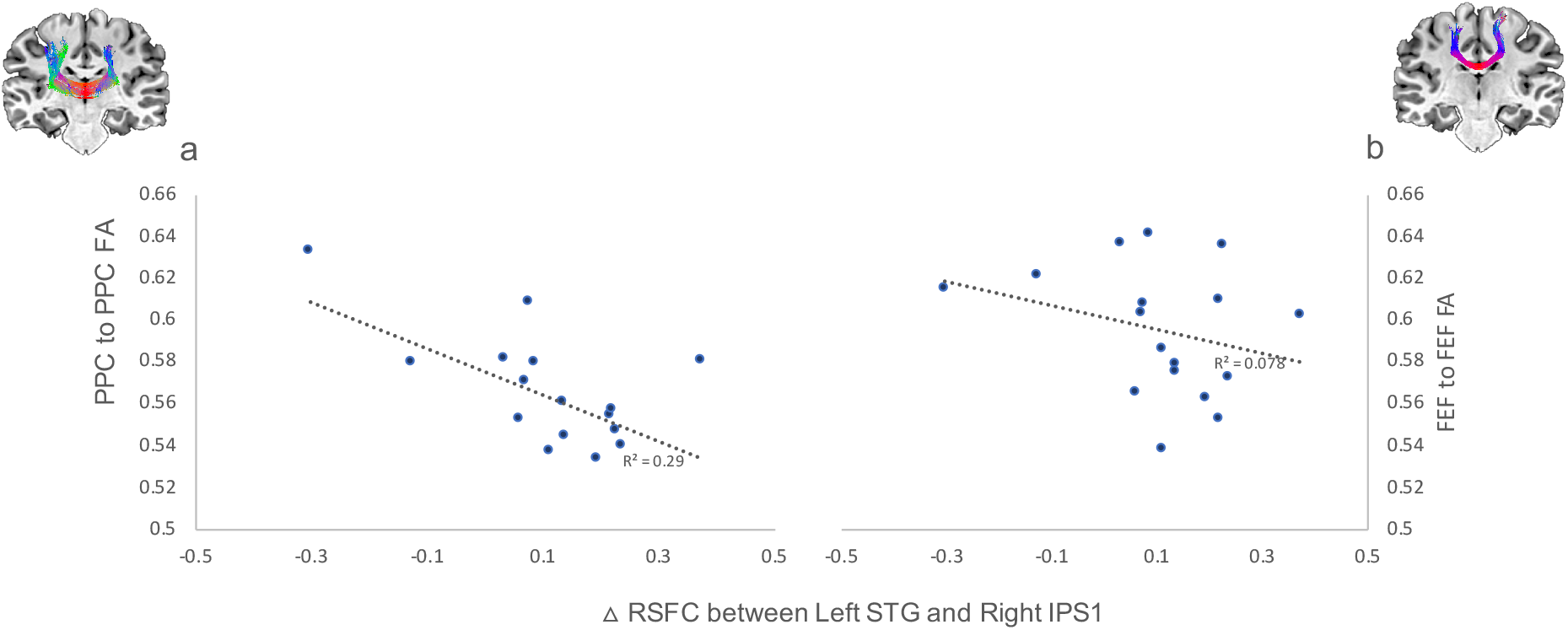
Association between the amount of change in RSFC between the left STG and right IPS1/2 seed after TMS and callosal FA between **a)** left and right PPC and **b)** FEF, across participants.

## 4. DISCUSSION

We hypothesized that inhibitory TMS delivered to the right PPC of healthy individuals would produce a rightward visuospatial bias, decrease RSFC between the stimulated right PPC and frontal areas and increase RSFC with areas of the contralateral hemisphere. We also expected that structural connectivity between the left and right PPC would predict the magnitude of RSFC changes in the left hemisphere.

As expected, continuous theta burst TMS of the right PPC produced a rightward visuospatial bias, as measured by the perceptual line bisection task. This finding reproduces previously reported results (e.g., Fierro et al., 2000; Szczepanski and Kastner, 2013) and, since the behavioral data were collected at the end of the experiment, shows that the TMS-induced effect was active during the fMRI acquisition. We also found a rightward shift in the subjective perceived midline on the straight-ahead pointing task. However, this shift was only present at the early-post measurement. The direction of change is consistent with inhibition of the right PPC and the trend is visible in Figure 4c. However, it was not borne out by a significant Time x Stimulation Location interaction. The absence of a statistically significant change on this behavioral measure is not surprising, since feedback from the motor performance requirement of the task likely reversed the visuospatial modulation (Schintu et al., 2020a).

Seed-based analysis of RSFC at the whole-brain level showed that TMS of the right PPC increased its RSFC with the left STG. The STG has been cited, along with the PPC (Critchley, 1953) and inferior parietal lobule (IPL; Vallar and Perani, 1986; Mort et al., 2003), as the source of spatial neglect when lesioned in both monkeys and humans (Karnath et al., 2001). The STG is generally considered part of the ventral attentional network (Shulman et al., 2010), which generates the non-spatial component of attention. However, STG activity also influences the dorsal network, which provides the spatial component (Corbetta and Shulman, 2011). Support for STG involvement in spatial attention comes from neglect patients in whom impaired functional connectivity between STG and middle frontal gyrus (MFG) has been shown to correlate with the impaired interhemispheric parietal connectivity, which is in turn associated with the severity of the attentional deficit (He et al. 2007). Consistent with lesion studies (Verdon et al., 2010) and both fMRI and TMS in healthy individuals (Galati et al., 2000; Neggers et al., 2006; Shah-Basak et al., 2018), STG inhibition by TMS affects allocentric, object-based, spatial processing (Ellison et al., 2004; Shah-Basak et al., 2018). It is noteworthy that the left STG cluster which increased RSFC with the right PPC seed is close to Heschl’s gyrus, where early cortical auditory processing occurs (Tzourio et al., 1997). However, this change in RSFC cannot be attributed to the TMS auditory artifact (Siebner et al., 1999), since the connectivity change was unilateral whereas the auditory artifact is binaural, and the direction of change was not identical in both PPC and vertex conditions (for which the auditory artifact is comparable).

As expected, we found a change in interhemispheric RSFC, but not between the PPCs, as we predicted based on previous studies. This is surprising if framed by the rivalry theory and the re-activation of the right PPC and de-activation of the left in recovery from neglect (Corbetta et al., 2005). However, a previous study (Ricci et al., 2012) applying TMS over the right PPC of healthy participants found a bilateral decrease in PPC activation, as though the PPCs were being modulated in parallel. This apparent inconsistency can be explained by context dependency. Blankenburg et al. (2008), in a concurrent TMS/fMRI experiment, found that TMS over the right parietal cortex enhanced activity in the left somatosensory area evoked by right median nerve stimulation, but had the opposite effect at rest. Another instance of the two PPCs behaving as a unit comes from a recent study from our laboratory (Schintu et al., 2020b), where we investigated the changes in RSFC after adaptation to left-shifting prisms, and found a bilateral decrease in RSFC within each of the two PPCs, along with the expected, neglect-like, leftward, shift in attentional bias previously described (Schintu et al., 2014). The existence of PPC to PPC interhemispheric inhibition has been shown in healthy individuals by online TMS effects on behavior (Szczepanski and Kastner, 2013) and electrophysiological measures (Koch et al., 2011). To date, there is no evidence of reciprocal inhibition between the PPCs at rest in healthy individuals. Based on these and our earlier results (Schintu et al., 2020b) it seems that rivalry between the PPCs in healthy individuals is evident only when PPC is preferentially activated either by TMS stimulation or task engagement. This is also true for the primary motor areas, which are also linked by inhibitory connections. (Ferbert et al., 1992). There, unilateral voluntary muscle contraction increases the interhemispheric inhibitory from the activated motor area to its contralateral homolog (Vercauteren et al., 2008). Another remote possibility for the absence of a change in RSFC between the two PPCs in the present finding, might be such modulation did not survive the cluster threshold despite the power calculation.

The reshaping of the attentional network following temporary disruption of the right PPC is substantiated by the correlation between PPC to PPC white matter FA and the magnitude of RSFC change across participants. The negative correlation implies that stronger structural connectivity between the two PPCs protects against TMS-induce disruption. Furthermore, the fact that the PPC to PPC FA predicted the increase in right PPC to left STG RSFC supports the idea that the transcallosal connection allows two PPCs to act as a unit, at least at rest, and split under activity.

In conclusion, we have shown that inhibition of the right PPC affects causes reorganization of the attentional network, detectable at rest, and that this change is mediated by the callosal connection between the PPCs. These findings also underscore the integral role of the STG in the functioning of the attentional network.

### CRediT AUTHORSHIP CONTRIBUTION STATEMENT

**Selene Schintu:** Conceptualization, Investigation, Formal analysis, Writing – original draft, Funding acquisition. **Catherine A. Cunningham:** Investigation, Formal analysis. **Michael Freedberg:** Methodology, Writing – review & editing. **Paul Taylor:** Supervision, Methodology. **Stephen J. Gotts:** Supervision, Writing – review & editing. **Sarah Shomstein:** Conceptualization, Writing – review & editing, Funding acquisition. **Eric M. Wassermann:** Conceptualization, Writing – original draft, review & editing, Funding acquisition.

## DECLARATION OF COMPETING INTEREST

The authors declare no conflict of interest.

## ACKNOWLEDGEMENTS

We would like to thank Joelle Sarlls, PhD and Vinai Roopchansingh, PhD for their support in designing MRI sequences.

## FUNDING SOURCES

This work was supported by the National Institutes of Health Ruth L. Kirschstein National Research Service Award to Dr. Schintu, the National Institute of Neurological Disorders and Stroke (1ZIANS002977-20) to Drs. Schintu, Wassermann and Freedberg, the National Science Foundation (grants BCS-1534823 and BCS-1921415) to Prof. Shomstein, the Center for Neuroscience and Regenerative Medicine (CNRM-70-3904) to Dr. Freedberg, and the Intramural Research Programs of the National Institute of Mental Health to Dr. Gotts.

## Notes

### Competing Interest Statement

The authors have declared no competing interest.

